# The spruce bark beetle *Ips typographus* transmits mutualistic fungi in mandibular mycetangia

**DOI:** 10.64898/2026.02.24.707679

**Authors:** Maximilian Lehenberger, Veit Grabe, Dineshkumar Kandasamy, Niklas Gentsch, Christoph Au, Ana Patricia Baños Quintana, Martin Schebeck, Martin Kaltenpoth, Jonathan Gershenzon

**Affiliations:** Department of Biochemistry, Max Planck Institute for Chemical Ecology, Jena, Germany; Microscopic Imaging Service Group, Max Planck Institute for Chemical Ecology, Jena, Germany; Department of Biology, Lund University, Lund, Sweden; Department of Insect Symbiosis, Max Planck Institute for Chemical Ecology, Jena, Germany; BOKU University, Department of Ecosystem Management, Climate and Biodiversity, Institute of Forest Entomology, Forest Pathology and Forest Protection, Vienna, Austria

**Keywords:** Bark beetle, *Ips*, symbioses, mycetangia, mutualist, PAS, µCT, *Picea abies*, fungi

## Abstract

Insects associated with mutualistic microbes often possess structures specialized for microbial transport that ensure the maintenance of these symbioses. Called mycetangia, these structures have evolved in numerous bark beetle species facilitating fungal transport to new host trees. The Eurasian spruce bark beetle *Ips typographus* is associated with several filamentous ascomycetes that may provide beetles with nutrition or help overcome tree defenses. *I. typographus* has been believed to vector its fungal symbionts mostly on the exoskeleton, but exact mechanisms of fungal transmission are unknown. Here, we report the discovery of a mandibular mycetangium in *I. typographus* that enables the transmission of symbiotic fungi. Extensive fungal isolation from *I. typographus* heads suggested the presence of a mycetangium while micro-computed tomography and histological analyses revealed a mycetangium close to the mandibles. Strikingly, we identified these fungus-carrying structures in both sexes of *I. typographus* and in the congeneric species *I. acuminatus*, and *I. duplicatus*, although mandibular mycetangia have been thought to be rare in *Ips* bark beetles. Moreover, mycetangia in male beetles in general were hardly reported in bark beetles before. The occurrence of mycetangia in *Ips* species highlights the important role mutualistic fungi play in the natural history of these bark beetles.

## Introduction

Bark beetles (Coleoptera: Curculionidae), a highly diverse group comprising more than 6,300 species worldwide, colonize various broadleaf and coniferous trees (1). While the majority of bark beetle species attack weakened or recently deceased trees, a few species consistently attack healthy trees (1, 2), causing extensive tree mortality in forests worldwide (2, 3). In general, two ecological strategies can be distinguished among bark beetles (*sensu lato*): Sapwood is colonized by fungus-farming ambrosia beetles, while phloem is colonized by the typical bark beetles (1, 3). Phloem-colonizing bark beetles drill tunnels through the protective outer bark of trees to get access to the energy-rich phloem (1) where they excavate large galleries for mating, reproduction, and feeding (4).

The phloem tissue of conifers is a challenging environment for bark beetles and other insects as it is chemically and structurally well-defended (5) and contains only low concentrations of many nutrients required for successful insect development, even though the phloem is enriched with simple sugars compared to the surrounding tissues (5–8). Moreover, the most abundant carbon sources, such as cellulose and hemicelluloses, are difficult to digest for most insects (7). Further, other nutrients such as B vitamins, sterols and elements such as phosphorous, potassium, nitrogen, and sulfur are severely limited (8–10). Consequently, many tree-colonizing insects benefit from symbiotic microbes, which can break down plant polysaccharides as well as polyphenolics and thus make the carbon and energy accessible to the insect host (7, 8, 11).

Many bark beetles as well as ambrosia beetles are well-known for their association with free-living, mutualistic fungi that may contribute to their survival in several ways (1, 2). First, mutualistic fungi can improve the nutritional quality of the substrate by providing various essential nutrients or by depolymerizing plant polysaccharides (8, 12–16). Second, they can help overcome the defenses of the host trees by metabolizing constitutive defense compounds and reducing the formation of induced defenses (5, 17, 18). Third, they may outcompete microbes that are pathogenic to beetles (19–21). To realize these benefits, bark beetles need to ensure the presence of mutualistic fungi in their galleries.

To maintain their association with mutualistic fungi, bark beetles and other insects have mechanisms for the transmission of these microbes across generations (22). Often, this involves the evolution of structures that are specialized for fungal transmission (23). For example, ambrosia beetles possess specialized reservoirs known as mycetangia (or mycangia) to ensure the transport of fungal symbionts to new host trees, (24–27). Within mycetangia, which have evolved convergently multiple times in ambrosia beetles (2), fungi are transported either as spores or as yeast-like cells (e.g. ambrosial growth) (28). Bark beetles such as *Dendroctonus* spp. possess mycetangia as well (29, 30), but these structures have been sought without success in several other species (31–34), suggesting different symbiont-transmission mechanisms or the occurrence of mycetangia that are difficult to detect.

The bark beetle species that has caused the most damage in European forests in recent years is the Eurasian spruce bark beetle *Ips typographus*, which mainly attacks Norway spruce (*Picea abies*) trees (3). Although weakened trees with reduced defense are typically colonized by *I. typographus* beetles, mass attacks on healthy trees can occur after abiotic disturbances such as storms and droughts, which can weaken tree defenses by decreasing resin flow and reducing carbon storages used for active defense responses (3, 35). As with other bark beetle species, the fungal symbionts of *I. typographus* are believed to promote beetle colonization of the host tree (5, 17, 36, 37). The ascomycetes *Grosmannia penicillata*, *Endoconidiophora polonica*, and *Ophiostoma bicolor* are considered the major symbionts (38–40), and *Ceratocystiopsis minuta* and *Ophiostoma brunneolum* are frequently isolated from *I. typographus* as well (41–43). As the fungal community of *I. typographus* is consistently found in freshly-established galleries, they are likely to be vectored by adult beetles. However, no mycetangia have been described for *I. typographus* so far and this beetle is considered to vector symbiotic microbes on their elytra or other body surfaces in cuticular pits (36). The lack of described mycetangia suggested that fungi are not essential for the successful colonization and development of *I. typographus* in host trees (44, 45). Further information on the mode of fungal symbiont transmission in this species is also essential to understand the maintenance and stability of its association with specific fungi.

Here, we report the discovery of paired mycetangia in *I. typographus*, which appear to ensure the successful transmission of fungal symbionts by adult beetles, as in other bark beetle species (28, 46, 47). First, we isolated filamentous fungi from the surface as well as from homogenized beetle heads from both sexes of *I. typographus* to identify the dominant taxa. Next, we utilized histological and x-ray approaches to search for fungal-carrying structures in *I. typographus* in comparison with other *Ips* species. Our analyses revealed the presence of a pair of small mandibular mycetangia in males and females of one of the most destructive forest pests in Europe, highlighting the importance of fungal symbionts in its success.

## Materials and Methods

### Study specimens

Dispersing adults of *Ips typographus* were collected during May/June 2024 using pheromone-baited traps (Pheroprax® and Ipsowit®, Witasek, Germany) in Norway spruce (*Picea abies*) stands near Tharandt (Germany) for almost all experiments. The SEM analyses employed beetles from an *I. typographus* culture reared on spruce logs in Lund, Sweden (Lund University). In general, traps were emptied twice a day and beetles were subsequently stored in petri dishes equipped with moist tissue and shipped to Jena at 4°C. Here, beetles were either stored at 4°C (for a maximum of 4 days) or immediately processed. The sex of all *I. typographus* individuals was determined by dissecting the abdomen and evaluating the reproductive organs. *Ips acuminatus* was obtained from naturally colonized trees of Scots pine, *Pinus sylvestris*, in northeastern Austria (Schönberg am Kamp) in spring 2024 that contained post-hibernation beetles from the winter season 2023/24 prior to emergence. Trees with signs of infestation, i.e. boring holes, were harvested and the trunk cut into pieces with a length of 60 cm. These small logs were transferred to incubators with 25 °C and 16 h light/8 h dark (BOKU University, Vienna) and checked for newly emerged beetles every day. Males and females were distinguished by morphological traits (i.e. spines on the elytral declivity) and express-mailed to Jena/Lund for subsequent analyses. *Ips duplicatus* from the Czech Republic, included in our µCT analyses, were sent alive to Jena and immediately processed.

### Isolation and identification of filamentous fungi

Filamentous fungi were isolated from the surface of adult *I. typographus* beetles as well as from homogenized heads to identify internally transmitted fungi. Overall, we isolated fungi from 18 females and 12 males. For the isolation of fungi from the beetle surface, each individual beetle was first placed in 500 µl of sterilized 1×PBS buffer with 0.1% Tween 20 and vortexed for approx. 5 s to wash off the majority of external fungi. Afterwards, beetles were transferred to a sterilized piece of aluminum foil using sterile forceps while constantly working under a Bunsen burner to create a sterile working environment. Heads were removed using sterile forceps and a scalpel, and were individually transferred to 500 µl of 1×PBS with 0.1% Tween 20, while the remaining abdomens were used to sex beetles via dissection of their reproductive organs. To homogenize beetle heads, we used a melted pipette tip followed by vortexing for approx. 15 s. The suspensions obtained were diluted 1:10 to reduce the abundance of fast-growing yeasts. Afterwards, the two suspensions from each individual beetle (surface and head) were inoculated in duplicate on potato-dextrose agar (PDA, Roth, Germany) as well as in triplicate on spruce phloem medium (2% freshly prepared, freeze-dried phloem powder, 2% agar from Roth, Germany) as a more selective growth substrate. We used two different types of media to optimize fungal isolations and to particularly search for slow growing fungi, which might otherwise be overgrown. On each petri dish and medium, 100 µl of one of the suspensions were streaked out and incubated at 25 °C and 65% humidity. Plates were checked daily and filamentous fungi were isolated from each plate. Pure cultures from each isolate were obtained after another sub-culturing on PDA and were divided into different morphotypes. Finally, cryo stocks (glycerol / water: 60 / 40) were prepared and stored at −80°C.

For the molecular identification, DNA was extracted from homogenized fungal tissue from all fungal morphotypes isolated from each individual beetle following the Qiagen DNeasy® Blood & Tissue Kit. For the PCR, we either amplified the internal transcribed spacer (ITS) or the large subunit (LSU) rRNA using the 2× MyTag Red Mix (Meridian) as master mix and 1 µl template (1:10 diluted). We used the general fungal primers ITS-1 (TCCGTAGGTGAACCTGCGG) and ITS-4 (TCCTCCGCTTATTGATATGC) for the ITS region of the ribosomal DNA (57) as well as the primer pair LROR (GTACCCGCTGAACTTAAGC) and LR-5 (ATCCTGAGGGAAACTTC) for the LSU region (58). Primers were purchased from Eurofins Genomics (Ebersberg, Germany). The following PCR conditions were applied: 95 °C for 1 min, followed by 35 cycles of 95 °C for 15 s, 54 °C (for the ITS Primers) or 56 °C (for the LSU primers) for 15 s, and 72 °C for 40 s, ending with 72 °C for 5 min and cooled down to 5 °C. After gel electrophoresis using 1% agarose gels (Roth, Germany) and the GelGreen®NucleicAcidStain (10.000 × water; Millipore, Germany), we used the Wizard® SV Gel and PCR Clean-up System (Promega, Germany).

Amplified ITS and LSU regions were then subjected to Sanger sequencing. Here, purified DNA was diluted to a final concentration of 25-30 ng/µl before being added to the sequencing PCR with a master mix consisting of 2 µl of Big Dye (ThermoFisher Scientific, Germany), 2 µl of BigDye Terminator 5X Sequencing Buffer (ThermoFisher Scientific, Germany), 1 µl primer, 14 µl of nuclease-free water (Promega, Germany), and 1 µl template. Amplification for sequencing was performed with an initial denaturation of 94°C for 5 min, followed by 35 cycles of 30 s at 94 °C, 30 s at 55 °C, 4 min at 60°C, and no final extension. Amplified products were loaded onto an ABI Prism® Gen-Analysator 3130xl 16-capilaries following the manufacturer’s instructions. Sequence quality was first checked using the SnapGeneViewer 3.2 (SnapGene software, from GSL Biotech; available at snapgene.com), while fungi were finally identified using BLASTn at NCBI, where the best hit based on query cover and percentage identity was used for annotation (59). The most frequently occurring fungi were visualized by bar plots using the statistical software R (version 4.2.1) applying the packages “ggplot2” (60) and “girdExtra” (61). Additionally, statistical differences were revealed in R using Fisher’s exact test including Bonferroni correction to account for multiple testing.

### Micro-computed tomography (µCT)

Analyses were performed with adult males and females of *I. typographus*, *I. acuminatus*, and *I. duplicatus* (two individuals per sex and species). Living beetles were first fixed in 4% paraformaldehyde (PFA, Roth, Germany) in 80% ethanol (Roth, Germany) for 2-3 days at room temperature. After two washing steps (each 1 h) with 80% ethanol on a shaker at room temperature, beetles were further dehydrated in absolute ethanol for two to three days at room temperature. The samples were then placed in a freshly prepared 1% methanolic iodine solution (Methanol: VWR, Germany; Iodine: Sigma, Germany) for 24 hours of contrasting. To remove the excess iodine, three washing steps (each 1 h) in denatured absolute ethanol and three further washing steps (each 1 h) in 100% ethanol were carried out. Finally, the beetles were critically-point-dried using a Leica CPD300 auto (Leica Microsystems, Wetzlar, Germany) with a standard program (100% stirring, slow CO_2_ influx with a delay of 60 minutes, 35 exchange cycles at a speed of 3, 100% slow gas out at medium heat). For µCT analyses, beetles were mounted in cut pipette tips according to their size and glued with UV-curing glue (Fotoplast Gel) to the metal holder of the µCT (Bruker SkyScan 1272, Billerica, Massachusetts, USA). The scans were carried out with the x-ray source running at 25 kV and 125 µA as well as 360° rotation at 0.2° steps, 1.5 µm pixel size, an exposure time of 519 ms and 4 times averaging. Reconstruction of the x-ray projections was executed in NRecon, which included correcting the misalignment, ring artifacts, and beam hardening. Final scans were visualized and manually segmented using Dragonfly software (Version 2022.2 Build 1409; Comet Technologies Canada Inc., Montreal, Canada; software available at https://www.theobjects.com/dragonfly).

### PAS-based histology of beetles and fungi

Periodic acid-Schiff staining (PAS) was performed with adult males and females of *I. typographus* (N = 3 per sex) and *I. acuminatus* (N = 3 per sex). *I. duplicatus* was not included in this analysis due to a lack of sufficient material. Beetles were decapitated and the abdomens were used to reliably sex the specimen by genital dissection. The heads were directly fixed in 4% PFA in PBS (Roth, Germany) for 48 h on a shaker at room temperature. After multiple washing steps in PBS, the samples were dehydrated in an ascending tertiary butanol (Roth, Germany) series (30%, 50%, 70%, 80%, 90%, 96%, three times 100%) for 2 h for each step on a shaker at room temperature. In preparation for the Histocure 8100 embedding, samples were transferred three times (each 2 h) to 100% acetone (Roth, Germany), followed by 1 h in acetone and infiltration solution in a 1:1 mix and finally three times (each 2 h) in pure infiltration solution of Histocure 8100 (Morphisto, Offenbach, Germany). Next, the specimens were transferred into the Histocure 8100 embedding solution in sample mounts, then cured overnight on an open lab bench and for another night in a desiccator. The cured blocks were cut into 5 µm sections on a HistoCore Autocut R microtome (Leica Microsystems GmbH, Wetzlar, Germany) and transferred to microscope slides. For staining, the slides were incubated in periodic acid (5 min; Roth, Germany), rinsed in water (10 min), immersed in the reduction solution (3 sec; Roth Germany), and incubated in Schiff’s solution (5 min; Roth, Germany). After staining, slides were incubated in disulfite water (1 min, 2x; Roth, Germany), rinsed in water (10 min), counter-stained in 0.2% Fast Green dye (2 min; Omikron, Rietberg, Germany), rinsed in water (2 min), dehydrated in 96% ethanol (3 sec), rinsed in water (3 sec) and dried at 60 °C. After briefly submerging the dry samples in Xylol (Roth, Germany) and drying them again, they were mounted in Histokitt medium (Carl Roth GmbH + Co. KG, Karlsruhe, Germany) and covered with a glass slide for inspection under a DMi8 Thunder microscope (Leica Microsystems GmbH, Wetzlar, Germany) equipped with an HC PL APO 20x/0.80 objective for overview and a HC PL APO 40x/0.95 objective for detail views. Images were acquired using a Flexacam K3C.

### Scanning electron microscopy (SEM)

Adult male and female *I. typographus* (N = 3 per sex) were morphologically sexed prior to the analyses, air-dried for at least 10 days and mounted on metal stubs. To facilitate visualization of the interior of mycetangia, mandibles were detached from the mouth parts of selected samples using fine forceps. The samples were then sputter-coated with gold using a Cressington 108 auto (Cressington, Watford, UK) for 75 seconds at 20 mA. Mycetangia were examined with a Hitachi SU3500 SEM operated at 5 kV.

## Results and Discussion

To determine whether the fungal symbionts of *I. typographus* are predominantly carried on the beetle’s cuticle or within a mycetangium, we first applied a culture-dependent approach. We used two different types of artificial medium to isolate filamentous fungi from washes of the beetle’s body surface and from individual homogenized beetle heads after surface-washing. After generating pure fungal cultures, species were identified applying Sanger sequencing of the LSU or ITS region. Overall, we isolated 34 different filamentous fungi from both surface washes and heads of adult *I. typographus* (Suppl. Table S1). *Grosmannia penicillata*, C*eratocystiopsis minuta*, *Ophiostoma brunneolum*, *Aureobasidium pullulans*, and *Ophiostoma bicolor* were among the most common species and were generally isolated more frequently from heads than from surface washes for both sexes, except for *A. pullulans* and *O. brunneolum* in males (see Fig. 1). All of these are known *I. typographus* associates, often believed to be mutualists (36, 37, 41–43, 48, 49), except for *A. pullulans*, which is a ubiquitous and widespread saprophyte (50). In females, *G. penicillata*, *C. minuta*, and *O. brunneolum* were found more often in heads (occurring in 72.2%, 44.4% and 66.7% of female heads, respectively) compared to surface washes (occurring in 11.1%, 22.2%, and 27.8%, respectively; statistically significant for *G. penicillata*, Suppl. Table S2). The fungi *A. pullulans* and *O. bicolor* were isolated in lower frequencies from both sources. In males, *G. penicillata*, *C. minuta*, and *O. bicolor* were isolated more often from heads (occurring in 33.3%, 66.7% and 41.7% of male heads, respectively) and were less prevalent in the surface wash (occurring in 8.3%, 25%, and 16.7%, respectively). Meanwhile, *O. brunneolum* and *A. pullulans* were especially dominant in the surface wash (58.3%, and 66.7%, respectively), but were also often isolated from heads (58.3% and 33.3%, respectively).

**Figure 1:**
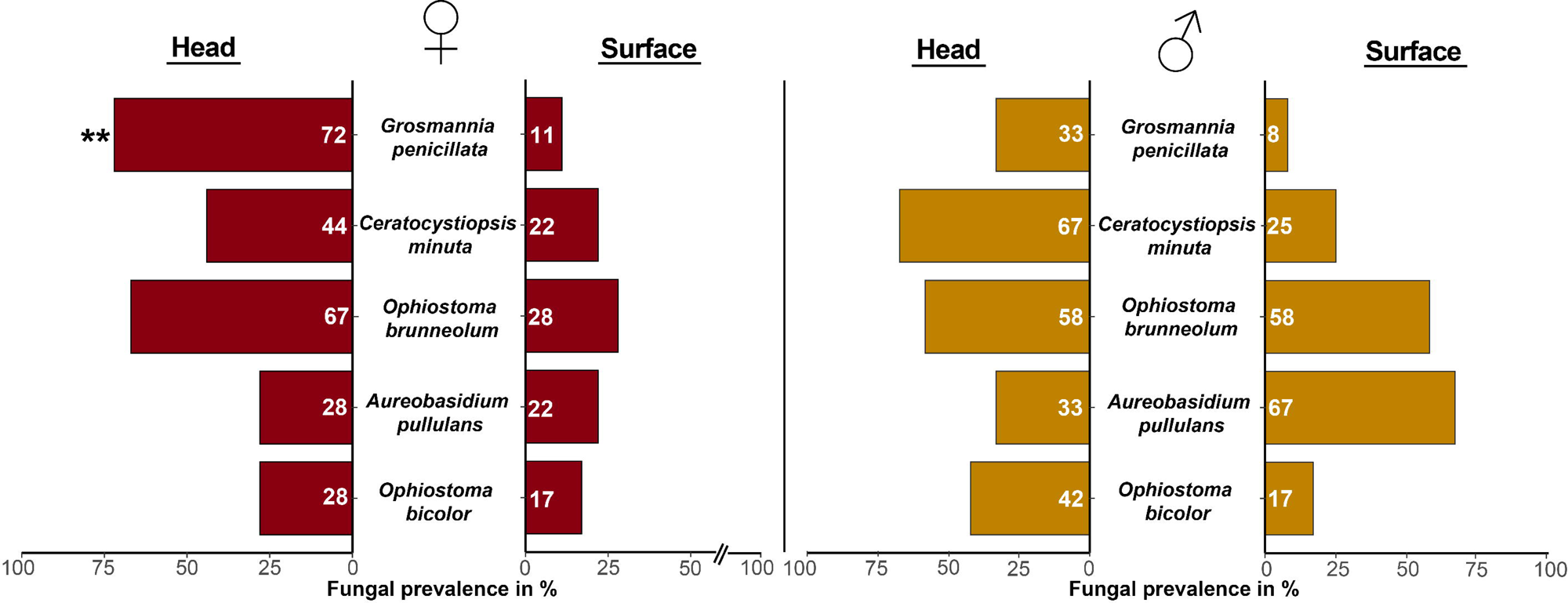
*Ips typographus* transports fungal symbionts more frequently in heads than on the body surface, consistent with a specialized head structure (= mycetangium) for fungal transmission. Bar plots show the most prevalent fungal taxa isolated from adult *I. typographus*. Samples were taken from crushed heads or body surfaces of 18 females (red bars) and 12 males (orange bars). X-axis shows the prevalence of each species in head and surface collections in percent, and the numbers in bars denote the exact percentage. Statistical differences were determined using Fisher’s exact test including Bonferroni correction to account for multiple testing (ns - P > 0.05, * - P < 0.05, ** - P < 0.01, *** - P < 0.001; see Suppl. Table S2 for individual p-values). An overview of all isolated fungi is provided in Suppl. Table S1.

The higher prevalence of well-known *I. typographus* associates in heads in comparison to the surface washes suggested the presence of a fungus-carrying mycetangium in the head. Thus, we performed micro-computed tomography (µCT) with field-collected female and male beetles. Here, we included a total of three different *Ips* species (*I. typographus*, *I. acuminatus,* and *I. duplicatus*) to enable the comparison of structural similarities within this genus. Interestingly, our µCT analyses revealed tiny paired mandibular structures between the base of the mandibles and the anterior tentorial arms in the ventral region of the beetle head, which were found in both sexes in all three *Ips* species (Fig. 2). These structures are similar to what had been described as mycetangia for *I. acuminatus* females before. However, our findings contradict the previous report that males of *I. acuminatus* lack mycetangia (25, 47). Indeed, mycetangia from bark and ambrosia beetles have almost exclusively been described from females (24, 25, 28, 47, 51). In this study, the size and dimensions of these structures were comparable between the three *Ips* species (including the different sexes) investigated with a length of the pouch ranging from 90 to 180 µm and a thickness ranging from 20 to 50 µm. Nevertheless, this mycetangium is much smaller than those described from other bark and ambrosia beetles (25, 52), indicating that the potential fungal load might be more limited.

**Figure 2:**
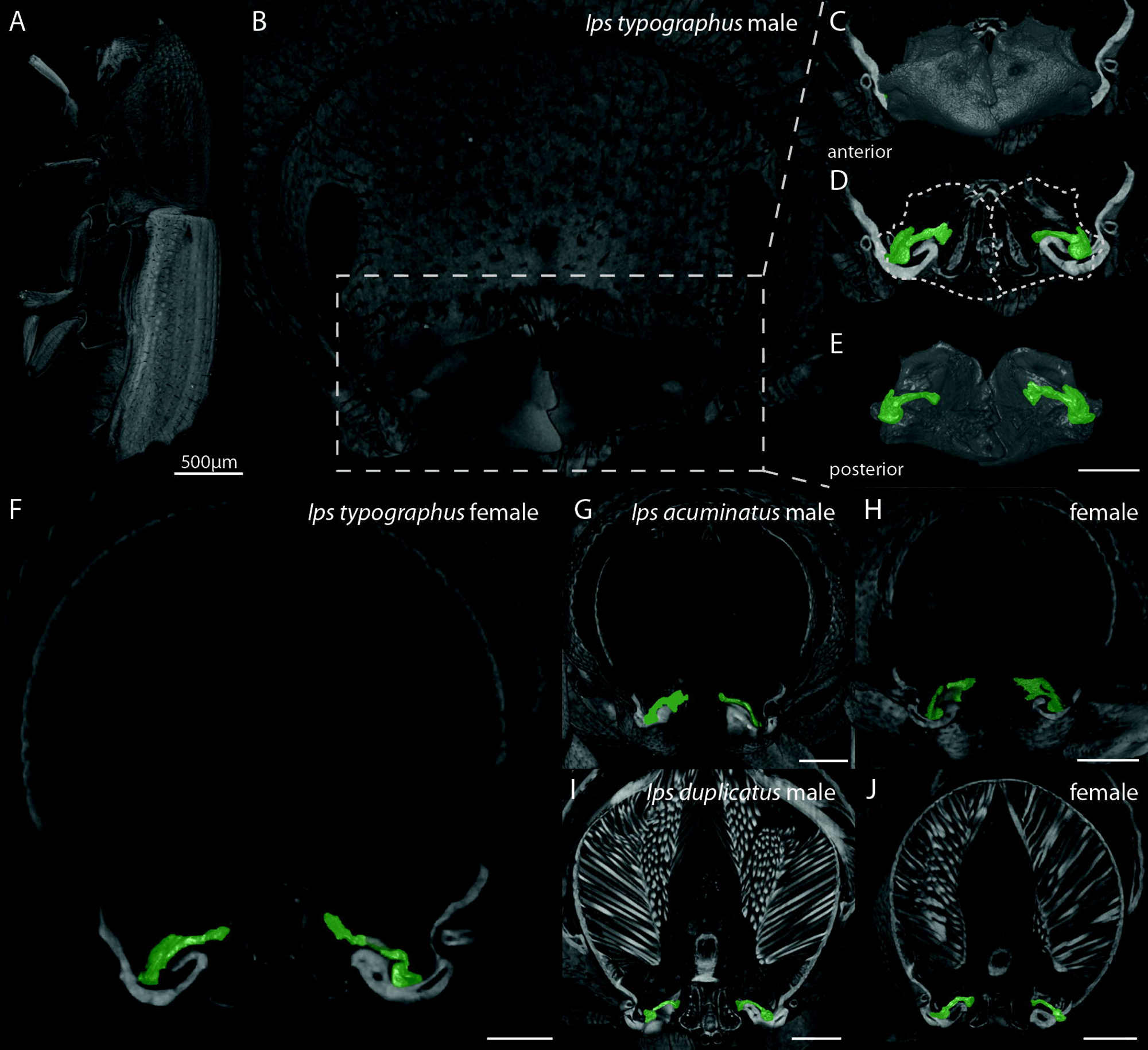
Micro-computed tomography (µCT) reveals paired mandibular structures in the heads of male and female *Ips typographus, I. acuminatus,* and *I. duplicatus*. (**A**) Lateral view of a male *I. typographus*. (**B**) Frontal view of the lower region of the head. (**C**) Frontal view of the mandible area (boxed in **B**) with mandibles in gray. (**D**) Frontal view with mandibles removed: Dashed white outline shows former location of mandibles; mycetangium in green. (**E**) Posterior view of the mandible area. (**F**) Frontal scan of a female *I. typographus* chosen to visualize the mycetangia. Identical perspectives are shown of the *I. acuminatus* male (**G**) and female (**H**); and the *I. duplicatus* male (**I**) and female (**J**). Scale bars indicate 200 µm if not stated otherwise.

As our µCT analyses suggested the presence of a mycetangium in the *Ips* species examined, we applied light microscopy to investigate the nature of this structure in detail. We treated the transverse head sections with periodic-acid Schiff staining (PAS) to confirm the presence of fungal cells, since fungal cell wall polysaccharides turn a characteristic purple or magenta color during this treatment (46, 53). The mycetangia of both sexes of *I. typographus* were indeed filled with fungi (Fig. 3 and Suppl. Fig. S1), which confirmed them as fungal symbiont-transmitting organs. PAS staining revealed approximately 12 individual fungal cells per mycetangium section. These quantities are much lower than the number of fungal cells typically found in other beetle mycetangia, such as for the ambrosia beetles *Trypodendron* spp. and *Xylosandrus germanus* (24). Curiously, in our *I. acuminatus* samples, the mycetangia of females (N = 3) were always filled with fungi, but those of males (N = 4) were always empty.

**Figure 3:**
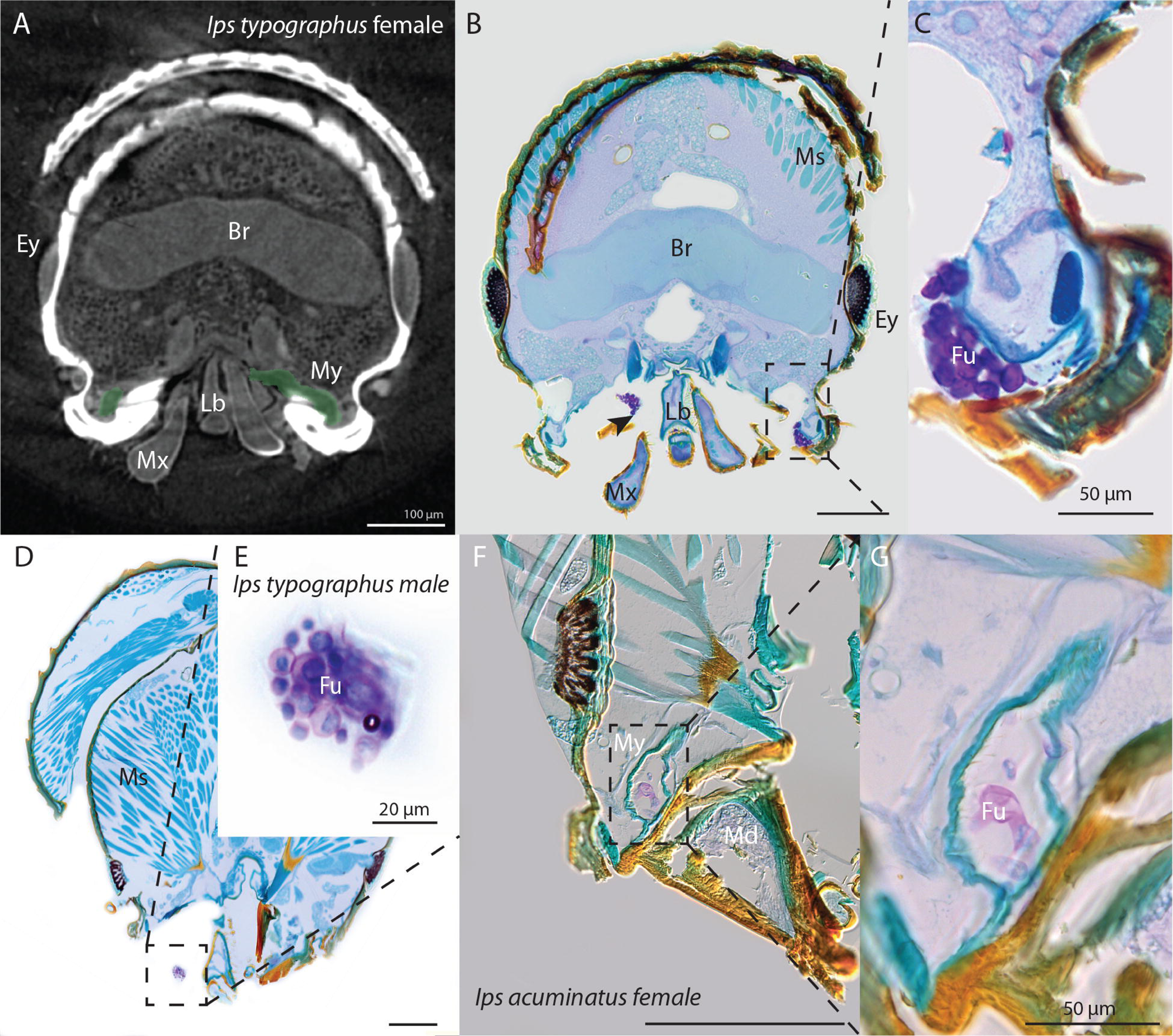
Light microscopy illustrates mycetangial structure and confirms the presence of fungal cells in both sexes of *I. typographus* and female *I. acuminatus*. (**A**) Section of a reconstructed µCT frontal scan of an *I. typographus* female (mycetangia - green). (**B**) Semi-thin section with a similar plane as in (**A**) treated with periodic acid-Schiff (PAS) staining: fungal spores appear purple. (**C**) Enlargement of view in panel **B**. (**D**) Head section and (**E**) enlargement of an *I. typographus* male. (**F**) Head section(**G**) enlargement of an *I. acuminatus* female (**F**, **G**). Br - brain, Ey - eye, Fu - fungus, Md - mandible, Ms - muscle, My - Mycetangium, Mx - maxilla, Lb - labium. Scale bars indicate 200 µm if not stated otherwise.

The round shape of the fungal cells in the PAS-stained mycetangium differs from the sausage-shaped asexual spores reported for *G. penicillata* (54). In fact, the morphology of bark and ambrosia beetle symbionts has previously been noted to change after they entered mycetangia from a mycelial form to a yeast-like state, the so-called ambrosial growth, which is believed to serve as a dormant growth form that may survive harsh environmental conditions (24, 28). To date, it is still unclear what triggers this morphological change in the mycetangium or the reversal to mycelial growth needed when beetles arrive at a new host tree and the fungi contact substrates suitable for growth (28).

The location of the *I. typographus* mycetangium near the mouth suggests that fungi are taken up directly during feeding. In fact, feeding is closely connected with the loading of the mycetangium in time and space. Freshly eclosed adults (newly hatched from pupae) need to fill their mycetangium since it is likely to be empty after the pupal phase and they are preparing for dispersal flight to a new host tree (28). Bark beetles such as *I. typographus* are known to feed extensively after eclosion to attain sexual maturity as well as to mature their flight muscles (55), using chemical cues to locate fungi (37, 49). This raises the question of how symbiotic fungi can enter such a mandibular mycetangium. Scanning electron microscopy (SEM) identified a tiny opening inside the mouth at the base of the mandibles (Fig. 4), which might function in the uptake as well as the release of fungi. Here, we observed multiple hair-like structures close to the putative entrance, which were also observed in other mycetangia-harboring beetles and may facilitate the uptake and storage of specific fungi (24, 25). These hairs might also secrete bioactive compounds that promote the selectivity of the mycetangium for specific fungal taxa (28). Our results demonstrate that the bark beetles *I. typographus*, *I. acuminatus*, and *I. duplicatus* all possess a mandibular mycetangium, which is present in both sexes. The occurrence of this structure in both males and females indicates the important function of symbiotic fungi for both sexes, although the differences in fungal community composition between sexes points to somewhat different roles. The dominant fungus isolated from male *I. typographus* heads was *C. minuta* (carried in 67% of the males, but only 44% of the females), while for female heads the dominant fungus was *G. penicillata* (carried in 72% of the females, but only 33% of the males). Since in *Ips* life cycles, the adult males are the pioneer sex on the host that initiates the galleries, we propose that *C. minuta* may be important for a male-associated role, such as assisting in mass attacks by producing attractive semiochemicals for beetles. Females arrive on the host tree after being attracted by male pheromones, mate and construct tunnels for oviposition. Thus, we propose that *G. penicillata*, their principal head-localized fungus, may be especially required for providing nutrition and metabolism of host defenses essential for larval growth (17, 56). The major head-localized fungi were occasionally isolated from elsewhere on the body surface, which might be due to spores or mycelia present around the mandibles or elsewhere on the surface outside the mycetangium. Other species such as *O. bicolor* and *O. brunneolum* were isolated from both surface and heads, suggesting mycetangial and non-mycetangial transmission.

**Figure 4:**
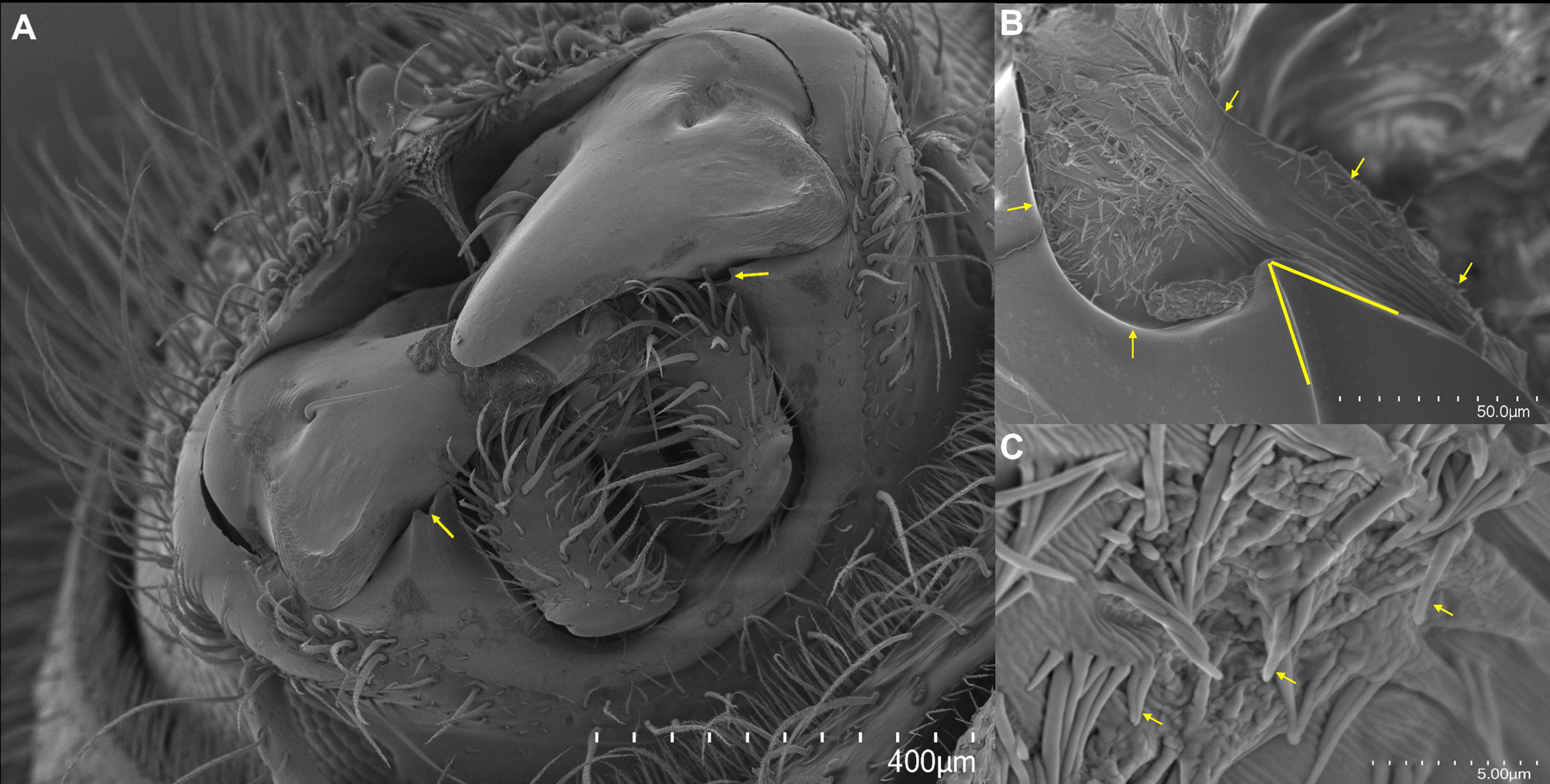
Scanning electron microscopy (SEM) shows potential entrances to the mandibular mycetangia in *Ips typographus*. Frontal view of the mouth area of a male *I. typographus*, where potential entrances to the mycetangia are indicated with white arrows on both mandibles. (**B**) Enlarged frontal view of the mandible area of a male *I. typographus* with mandibles removed to expose the inside of the mycetangium. Yellow arrows indicate the original position of the mycetangium. Yellow lines depict a possible connective channel which leads to the position of the entrance. (**C**) Enlarged view of the hair-like structures in a male *I. typographus* that are covering the internal surface of the mycetangia.

Prior to this study, both sexes of *I. typographus* and *I. duplicatus* as well as males of *I. acuminatus* were considered to transmit symbiotic fungi on their body surface without using highly developed morphological structures (36, 46, 47). This has led to the hypothesis that symbiotic fungi may not be critical for the success of these beetles. However, the evolution of a specialized mycetangium, despite its small size, indicates important roles of symbiotic fungi in these bark beetles and further strengthen the mutualistic relationship between such beetles and their fungal associates, similar to that in fungus-farming ambrosia beetles (1). Future work is needed to clarify which fungi are present in the mycangium and their specific functions in the life history of the beetle.

## Author contributions

ML, APBQ, and MS collected and sexted beetles. NG, CA, and ML isolated and sequenced fungi. VG and ML performed µCT analyses and PAS histology. DK performed SEM analyses. ML, MK, and JG designed the study and ML wrote the first draft. All authors revised the final version of the manuscript.

## Materials availability

This study did not generate new unique reagents.

## Data and code availability

All obtained fungal sequences reported herein are permanently archived at the National Center for Biotechnology Information (NCBI) with the accession numbers PQ897172-PQ897214 as well as PQ900724 - PQ900898.

## Acknowledgments

The authors would like to thank Natascha Rauch, Bettina Raquschke, and Domenica Schnabelrauch for their support in the laboratory and sequencing as well as Eva Papek, Elisabeth Ritzer, and Lara Kundtner for their help with the rearing of *Ips acuminatus* as well as Dr. Anna Jirosová for sending *Ips duplicatus* from the Czech Republic. Further, the authors are grateful to Ola Gustafsson at Lund University for his assistance with SEM analysis. Finally, the authors are thankful to Helene Francke-Grosmann for her preliminary work on the mycetangia of *Ips acuminatus*, which inspired this project.

