## Supplementary figures and images for "The spruce bark beetle *Ips typographus* transmits mutualistic fungi in mandibular mycetangia"

### Suppl. Fig. S1

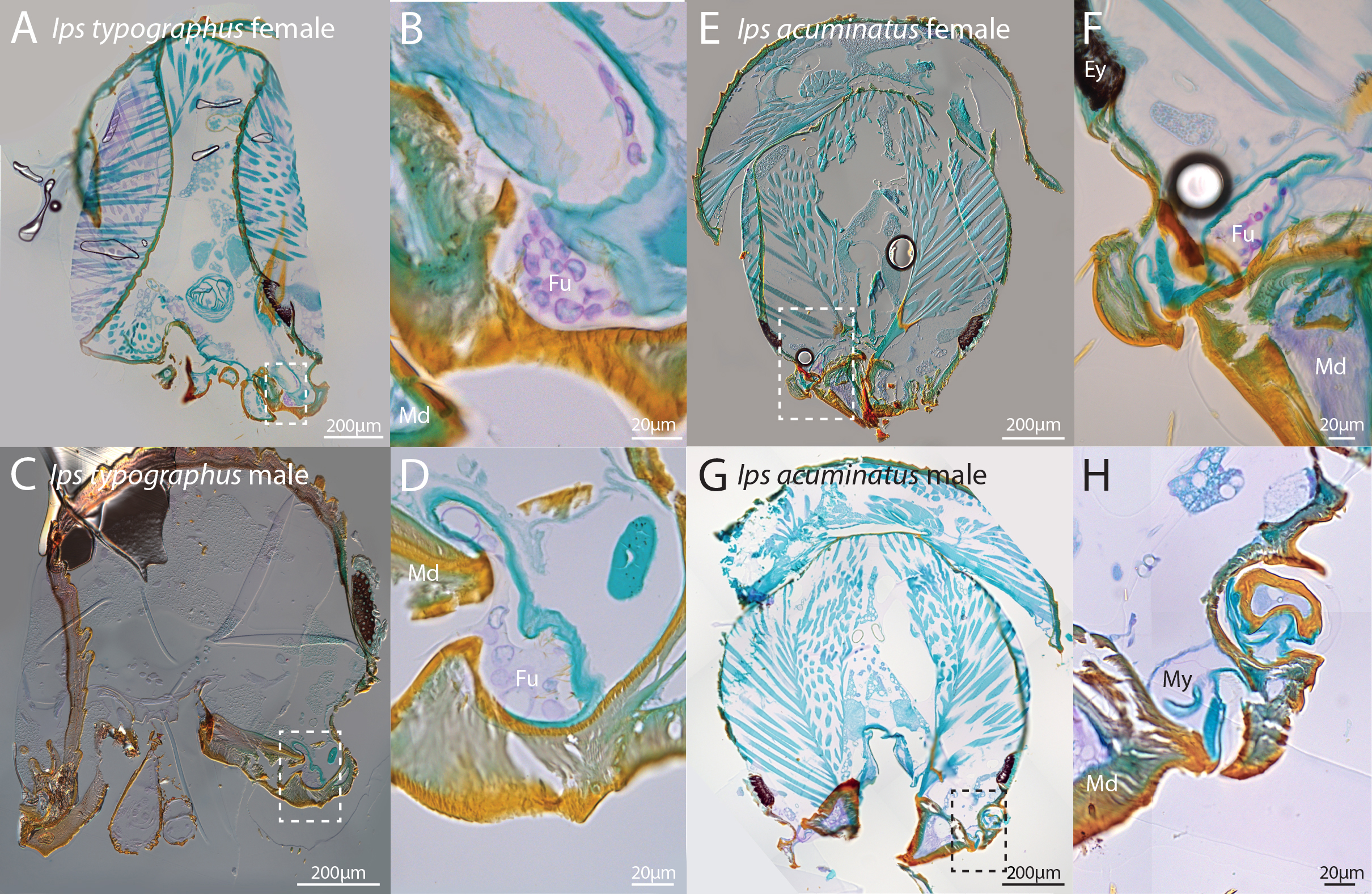
